# Integrated multi-omics investigations reveal the key role of synergistic microbial networks in removing plasticizer di-(2-ethylhexyl) phthalate from estuarine sediments

**DOI:** 10.1101/2021.03.24.436900

**Authors:** Sean Ting-Shyang Wei, Yi-Lung Chen, Yu-Wei Wu, Tien-Yu Wu, Yi-Li Lai, Po-Hsiang Wang, Wael Ismail, Tzong-Huei Lee, Yin-Ru Chiang

**Affiliations:** Biodiversity Research Center, Academia Sinica, Taipei, Taiwan; Department of Microbiology, Soochow University, Taipei, Taiwan; Graduate Institute of Biomedical Informatics, College of Medical Science and Technology, Taipei Medical University, Taipei, Taiwan; Institute of Environmental Engineering, National Central University, Taoyuan, Taiwan; Earth-Life Science Institute, Tokyo Institute of Technology, Tokyo, Japan; Environmental Biotechnology Program, Life Sciences Department, College of Graduate Studies, Arabian Gulf University, Manama, Bahrain; Institute of Fisheries Science, National Taiwan University, 106 Taipei, Taiwan

## Abstract

Di-(2-ethylhexyl) phthalate (DEHP) is the most widely used plasticizer worldwide with an annual global production of over eight million tons. Because of its improper disposal, endocrine-disrupting DEHP often accumulates in estuarine sediments in industrialized countries at sub-millimolar levels, resulting in adverse effects on both ecosystems and human beings. The microbial degraders and biodegradation pathways of DEHP in O_2_-limited estuarine sediments remain elusive. Here, we employed an integrated meta-omics approach to identify the DEHP degradation pathway and major degraders in this ecosystem. Estuarine sediments were treated with DEHP or its derived metabolites, *o*-phthalic acid and benzoic acid. The rate of DEHP degradation in denitrifying mesocosms was two times slower than that of *o*-phthalic acid, suggesting that side-chain hydrolysis of DEHP is the rate-limiting step of anaerobic DEHP degradation. On the basis of microbial community structures, functional gene expression, and metabolite profile analysis, we proposed that DEHP biodegradation in estuarine sediments is mainly achieved through synergistic networks between denitrifying proteobacteria. *Acidovorax* and *Sedimenticola* are the major degraders of DEHP side-chains; the resulting *o*-phthalic acid is mainly degraded by *Aestuariibacter* through the UbiD-dependent benzoyl-CoA pathway. We isolated and characterized *Acidovorax* sp. strain 210-6 and its extracellular hydrolase, which hydrolyzes both alkyl side-chains of DEHP. Interestingly, genes encoding DEHP/MEHP hydrolase and phthaloyl-CoA decarboxylase—key enzymes for side-chain hydrolysis and *o*-phthalic acid degradation, respectively—are flanked by transposases in these proteobacterial genomes, indicating that DEHP degradation capacity is likely transferred horizontally in microbial communities.

**Importance:** Xenobiotic phthalate esters (PAE) have been produced on a considerably large scale for only 70 years. The occurrence of endocrine-disrupting di-(2-ethylhexyl) phthalate (DEHP) in environments has raised public concern, and estuarine sediments are major DEHP reservoirs. Our multi-omics analyses indicated that complete DEHP degradation in O_2_-limited estuarine sediments depends on synergistic microbial networks between diverse denitrifying proteobacteria and uncultured candidates. Our data also suggest that the side-chain hydrolysis of DEHP, rather than *o*-phthalic acid activation, is the rate-limiting step in DEHP biodegradation within O_2_-limited estuarine sediments. Therefore, deciphering the bacterial ecophysiology and related biochemical mechanisms can help facilitate the practice of bioremediation in O_2_-limited environments. Furthermore, the DEHP hydrolase genes of active DEHP degraders can be used as molecular markers to monitor environmental DEHP degradation. Finally, future studies on the directed evolution of identified DEHP/MEHP hydrolase would bring a more catalytically efficient DEHP/MEHP hydrolase into practice.

## Introduction

Phthalate esters (PAEs) are commonly used as plasticizers to improve the flexibility of plastics and their adhesive capacity in some aqueous products [1, 2]. The annual production of PAEs has increased up to 8 million tons in 2011 [2]. The side-chains of PAEs are composed of linear or branched aliphatic alcohols with varying lengths [2, 3]. Di-(2-ethylhexyl) phthalate (DEHP), also named bis(2-ethylhexyl) phthalate), is the most widely used plasticizers worldwide, accounting for approximately one-third and 80% of PAEs produced in the European Union and China, respectively. Because of the daily use of plastic worldwide and the easily diffusion of plastic polymers into the environment [4], PAEs can be detected almost everywhere in industrialized countries [2] as well as in polar regions [5].

The endocrine-disrupting and carcinogenic activities of PAEs in higher animals have raised substantial public concern [4]. For example, some studies have reported that in some fishery species, embryo maturation and oogenesis are impeded by even a low concentration of DEHP—0.1–0.2 µg/L [7, 8]; the DEHP concentration in some freshwater ecosystems is up to a much higher 21 µg/L [2]. In addition, DEHP not only affects endocrine and nervous systems but is also genotoxic to humans [9, 10].

Abiotically, DEHP can be photodegraded in surface water or the atmosphere with a degradation half-life ranging from weeks to years [11, 12]. However, DEHP does not undergo photodegradation in aquatic sediments lacking sunlight exposure and O_2_ [2]. Moreover, the high hydrophobicity of DEHP (its water solubility is approximately 3 mg/L) results in the adsorption of DEHP onto aquatic sediment particles [13], meaning that the content of DEHP is considerably higher in aquatic sediments than in surface water. For example, in South Africa, the DEHP concentration is approximately 6 µg/L in river water, whereas it is up to 3,660 µg/kg in sediments in the same river [2]. Moreover, the salinity of estuarine environments enhances the adsorption of DEHP onto estuarine sediments because of salt fractionation [13].

Biodegradation is the primary process through which PAEs are removed in municipal wastewater treatment plants [2]. Several studies have reported aerobic microbial degradation pathways for different PAEs [3]. In the aerobic pathway, DEHP is initially transformed into *o*-phthalic acid and 2-ethylhexanol through mono-(2-ethylhexyl) phthalate (MEHP) by dialkyl phthalate hydrolase and monoalkyl phthalate hydrolase [14–17] (**Fig. 1**). Subsequently, *o*-phthalic acid is converted to protocatechuate (3,4-dihydroxybenzoate), which is followed by ring-cleavage through either the *ortho*-cleavage (by intradiol dioxygenases) or *meta*-cleavage (by extradiol dioxygenases) pathway (**Fig. 1**) [3,17,18]. By contrast, the anaerobic DEHP biodegradation pathway remains unclear. Nevertheless, the anaerobic degradation pathway of *o*-phthalic acid has recently been characterized in some denitrifying bacteria and sulfate-reducing bacteria [19, 20]. Briefly, *o*-phthalic acid is activated by either type III CoA transferase or ATP-dependent CoA transferase to form the highly unstable phthaloyl-CoA; this is followed by non-oxidative decarboxylation to form benzoyl-CoA by prenylated flavin-mononucleotide (FMN)-dependent phthaloyl-CoA decarboxylase in the UbiD family (21). Subsequently, the core-ring of benzoyl-CoA is cleaved through the well-established benzoyl-CoA degradation pathway [17,19,20] (**Fig. 1**). However, whether these *o-* phthalic acid-degrading anaerobes can degrade DEHP remains unclear.

**Figure 1.**
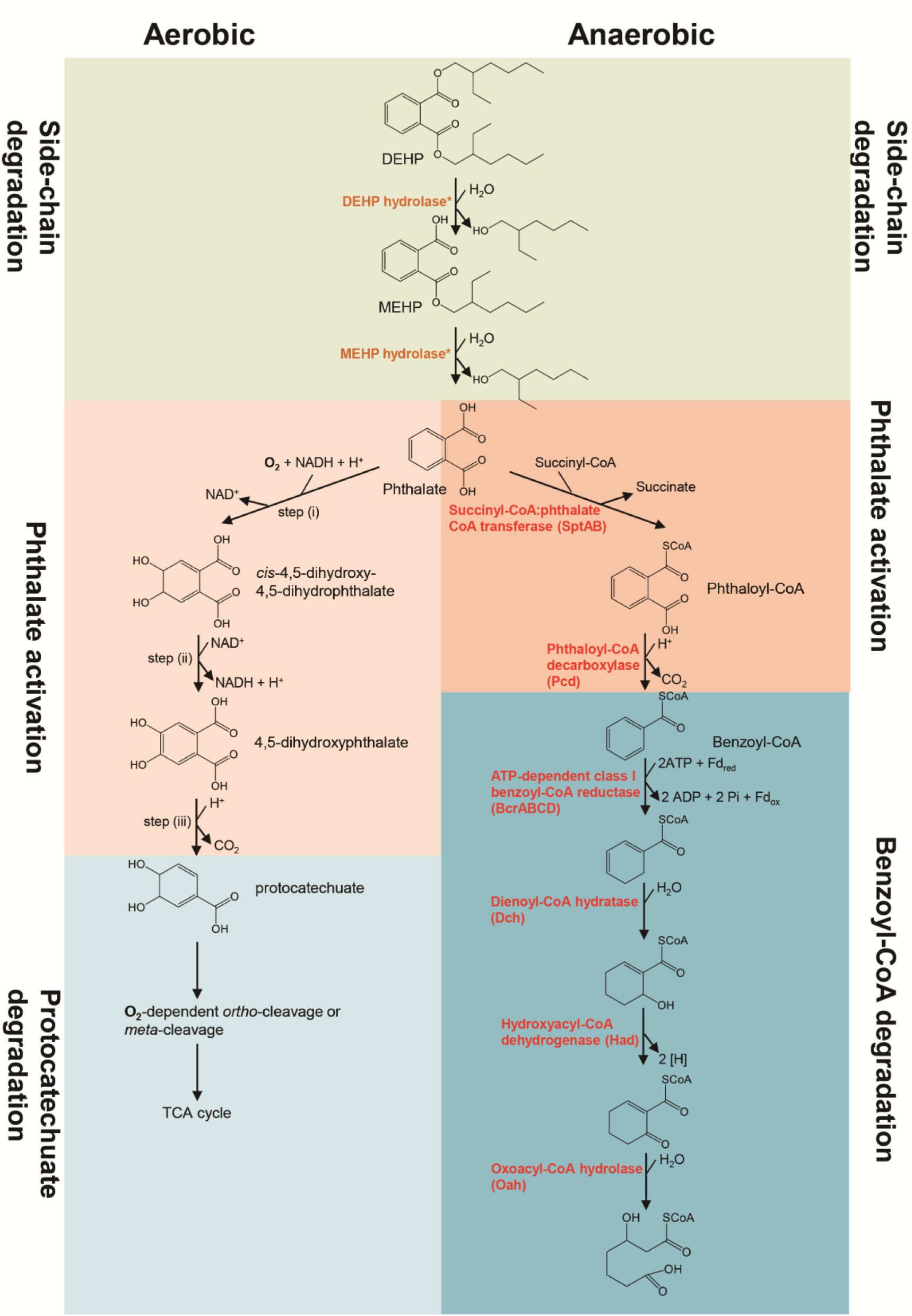
The proposed microbial degradation pathways of DEHP. Left panel: a simplified aerobic degradation pathway with 4,5-dihydroxyphthalate and protocatechuate as characteristic intermediates. An alternative aerobic pathway with 3,4-dihydroxyphthalate and protocatechuate as characteristic intermediates has also been identified (ref). Right panel: a proposed anaerobic degradation pathway for PAEs established in denitrifying *Aromatoleum*, *Azoarcus*, and *Thauera*. Proteins involved in this anaerobic pathway are shown in red.* Dialkyl phthalate and monoalkyl phthalate hydrolases were functionally characterized in aerobic *Gordonia* and anaerobic *Acidovorax* sp. strain 210-6 prior to this study. Fd, ferredoxin.

Wetland sediments in estuaries provide essential ecosystem services (water purification and toxin trapping) for urban environments and are major reservoirs of DEHP worldwide. To identify autochthonous anaerobic DEHP microbial degraders and elucidate the underlying biochemical and molecular mechanisms, we performed mesocosm experiments by incubating Guandu estuarine sediments with DEHP under denitrifying conditions. We used ultraperformance liquid chromatography (UPLC)–high-resolution mass spectrometry (HRMS) to identify DEHP-derived metabolites. Subsequently, we adopted next-generation sequencing approaches to identify DEHP-degrading bacteria and their degradation genes. Furthermore, we isolated the DEHP-degrading *Acidovorax* sp. and purified the corresponding DEHP/MEHP hydrolase from the sediment isolate. Both culture-independent and culture-dependent results suggested that DEHP was mainly removed from the estuarine sediments through synergistic microbial degradation, and side-chain hydrolysis represents a bottleneck in the DEHP degradation process.

## Results

### DEHP biodegradation in denitrifying sediment mesocosms

Sediment mesocosms (1 L; two replicates in each treatment) composed of estuarine sediment (approximately 200 g) and river water (approximately 800 mL) were treated with sodium nitrate alone [10 mM; abbreviated “SN (<u>s</u>ediment-<u>n</u>itrate)”], nitrate (10 mM) and benzoic acid [1mM; abbreviated “SBN (<u>s</u>ediment-<u>b</u>enzoic acid-<u>n</u>itrate)”], nitrate (10 mM) and *o*-phthalic acid [1mM; abbreviated “SPN (<u>s</u>ediment-*o*-<u>p</u>hthalic acid-<u>n</u>itrate)”], and nitrate (10 mM) and DEHP [1mM; abbreviated “SDN (<u>s</u>ediment-<u>D</u>EHP-<u>n</u>itrate)”]. Benzoic acid and *o*-phthalic acid have been proposed as crucial intermediates of the established anaerobic DEHP degradation pathway [17]. The concentrations of endogenous benzoic acid, *o*-phthalic acid, and DEHP in these sediment mesocosms ranged from 0.001 to 0.01 mM, approximately 100-fold lower than those in exogenous substrates. Sediment mesocosms that received different treatments displayed discernible patterns in substrate depletion rate and total nitrate consumption. The SN1 (nitrate alone; replicate 1) mesocosm had consumed 15.5 ± 0.92 mM nitrate in total after 25 days of anaerobic incubation (**Fig. 2AI**). In the SBN1 mesocosm, exogenous benzoic acid was largely depleted (approximately 80%) within four days of continuous nitrate consumption (**Fig. 2AII**). Approximately 80% of *o*-phthalic acid in the SPN1 mesocosm was consumed within one week (**Fig. 2AIII**). Notably, the DEHP consumption rate was lowest in the denitrifying sediment. Approximately 35% of the DEHP was consumed within 11 days, and the DEHP was depleted at day 21, together with a total nitrate consumption of 29.8 ± 0.16 mM (**Fig. 2AIV**). The mesocosms in all duplicates showed similar trends in the utilization of exogenous substrates (**Fig. S1**). We noted that the DEHP in the SDN2 was completely degraded at day 25, with the total nitrate consumption of 25.8 ± 0.22 mM of (**Fig S1D**).

**Figure 2.**
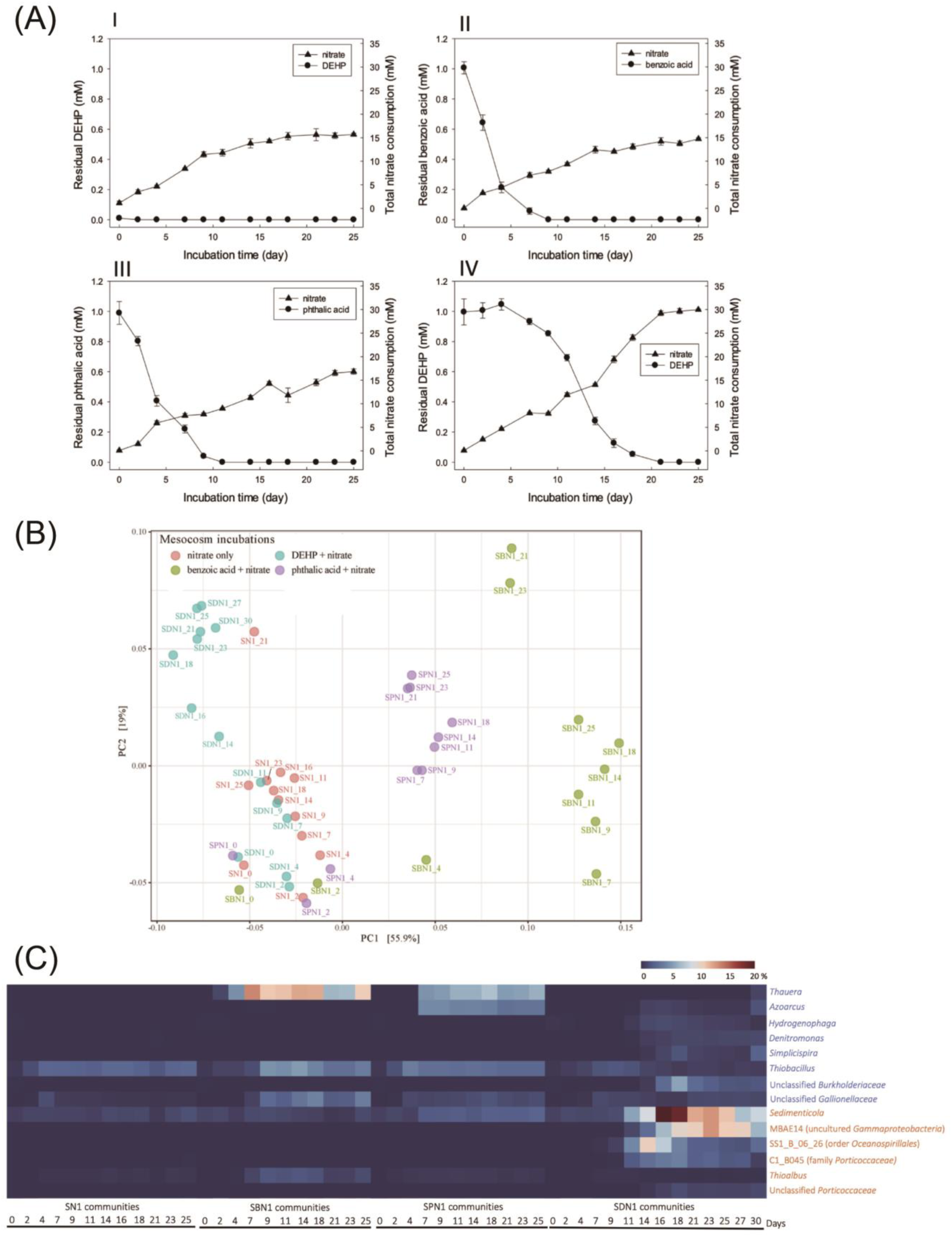
Estuarine sediments treated with benzoic acid, *o*-phthalic acid, and DEHP in denitrifying mesocosms. **(A)** Substrate utilization and nitrate consumption in (I) anoxic sediments incubated with nitrate only, (II) sediments incubated with benzoic acid and nitrate, (III) sediments incubated with *o*-phthalic acid and nitrate, and (IV) sediments incubated with DEHP and nitrate. The values are shown as means of three independent measurements with standard deviations for mesocosm replicate 1. **(B)** PCoA for the determination of similarities between the bacterial communities of mesocosm replicate 1 based on UniFrac distance matrix data (genus level) obtained from the sediment incubated with nitrate (SN1), sediment with benzoic acid and nitrate (SBN1), sediment with *o*-phthalic acid and nitrate (SPN1), and sediment with DEHP and nitrate (SDN1). **(C)** Relative abundance changes of genus in *Betaproteobacteria* (blue) and *Gammaproteobacteria* (orange) from sediment incubated with nitrate (SN1), with benzoic acid and nitrate (SBN1), with *o*-phthalic acid and nitrate (SPN1) and with DEHP and nitrate (SDN1) in mesocosm replicate 1. SS1_B_06_26 belongs to the order *Oceanospirillales* and C1_B045 belongs to the family *Porticoccaceae*.

DEHP-derived metabolites in the SDN1 samples were then identified through UPLC– Atmospheric Pressure Chemical Ionization (APCI)–HRMS (**Fig. 3**). We observed the production and subsequent consumption of two non-CoA thioester metabolites (MEHP and *o*-phthalic acid; **Fig. 3A**). These DEHP-derived metabolites were not detected within the first week of denitrifying incubation, whereas MEHP (0.08 ± 0.01 mM) and *o*-phthalic acid (0.19 ± 0.02 mM) had temporarily accumulated after 16 days of incubation (**Fig. 3B**).

**Figure 3.**
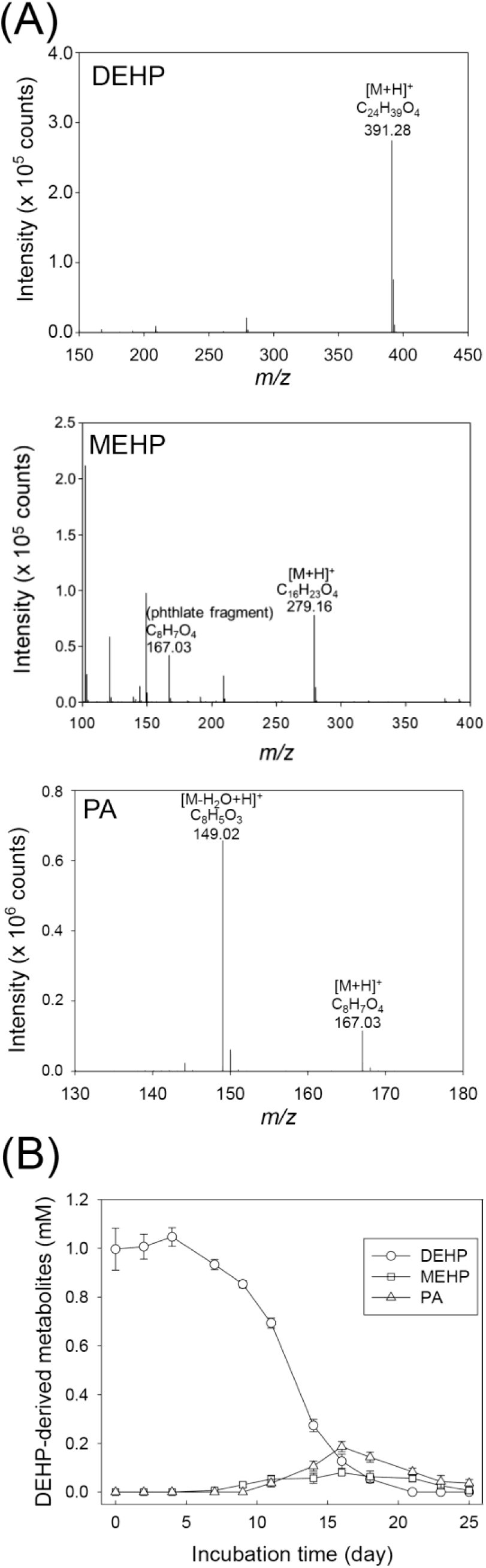
Metabolite profile analysis of DEHP-treated sediment mesocosm (replicate 1). (A) APCI–HRMS spectra of non-CoA metabolites detected in the mesocosm. PA, *o*-phthalic acid. (B) Time course of the DEHP consumption and the production of DEHP-derived metabolites in the sediment mesocosm. Quantification of the DEHP metabolites was based on pesudomolecular ([M+H]^+^) ion counts corresponding to individual compounds by using UPLC–HRMS. Data shown are the means ± standard deviations of three experimental measurements.

### Betaproteobacteria and gammaproteobacteria as key DEHP degraders in denitrifying sediment mesocosms

In total, 5,582 operational taxonomic units (OTUs) were generated in all the sediment communities of replicate 1. The PC1 (55.9% of total explained variance) and PC2 (19.0% of total explained variance) in the principal-coordinate analysis (PCoA) clearly distinguished the microbial communities among SN1, SBN1, SPN1, and SDN1. Most of the SN1 communities—together with communities from the early incubation stage (day 2 to day 4) of the SBN1, SPN1, and SDN1 mesocosms—were clustered in the lower-left corner of the coordinate, slightly distant from the sediment communities at the beginning of the experiment (incubation time of 30 min). However, communities from the middle and late incubation stages (day 9 to 30) of SBN1, SPN1, and SDN1 mesocosms were clustered separately (**Fig 2B**). The results of permutational multivariate analysis of variance (PERMANOVA) revealed that the weighted UniFrac distance of overall bacterial genera in each community was significant (F-value= 12.405, global R^2^= 0.45823, P-value <0.001).

In the benzoic acid and *o*-phthalic acid mesocosms, we observed enrichment of the betaproteobacteria *Thauera* and *Azoarcus* (**Fig. 2C**); this enrichment was associated with the consumption of the substrates within 1 week (**Fig. 2AII and 2AIII**). These two bacterial genera were not enriched in the SN1 mesocosm treated with nitrate alone. In the SBN1 community, the relative abundance of *Thauera* increased from <0.1% at day 0 to 12.5% at day 7. The abundance of *Thauera* and *Azoarcus* in the SPN1 community had increased to 5.8% and 3.9%, respectively, at day 11. Although *o*-phthalic acid was identified as a major DEHP metabolite in the SDN1 mesocosm (**Fig. 3**), the enrichment of the aforementioned two genera was considerably less (∼1%) in the SDN1 community. Instead, we observed large enrichment (≥10%) of three gammaproteobacterial genera—namely *Sedimenticola*, MBAE14 (uncultured *Gammaproteobacterium*), and SS1_B_06_26 (belonging to the order *Oceanospirillales*)—in the SDN1 community. In addition, the relative abundance of other bacteria, such as C1_B045 (gammaproteobacterial family *Porticoccaceae*) and an unclassified *Burkholderiaceae* (Betaproteobacteria), increased but to a lower extent (approximately 5%; **Fig. 2C**).

In the SN2, SBN2, SPN2, and SDN2 mesocosms (replicate 2), 5,708 OTUs were identified. The results of PCoA and temporal changes in the relative abundance of the aforementioned genera were similar to those observed in replicate 1 (**Figs. S2 & S3**).

### Putative carboxylesterase and β-oxidation genes involved in alkyl side-chain degradation in the DEHP-treated mesocosms

In differential gene expression (DGE) analysis, only 1,800 genes were observed to be up-regulated in the SPN sediments, whereas up to 40,867 genes were up-regulated in the SDN mesocosms (**Fig. S4**). In the SDN mesocosms, 205 alpha/beta hydrolase and esterase genes belonging to the carboxylesterase superfamily were annotated on the basis of eggNOG annotation in the DEHP-treated sediments. Only six genes–k141_3815848_2, k141_2812817_2, k141_4060503_4, k141_138169_2, k141_2938615_2, and k141_853245_1–were similar to the functionally characterized hydrolase gene NCU65476 of the denitrifying *Acidovorax* sp. strain 210-6 according to the hidden Markov model (HMM) search (the isolation and characterization of strain 210-6 and its DEHP/MEHP hydrolase are described later). Proteins encoded by these genes belonged to the IPR029058 carboxylesterase superfamily, and signal peptides were present on the proteins of k141_3815848_2, k141_2812817_2, k141_4060503_4, and k141_853245_1 **(Table 1).** After examining the binned genomes recovered from metagenomes (described in the next section), these genes were distributed on Bin6, Bin13, Bin14, Bin18, and Bin 44, and their taxonomic affiliations were *Acidovorax* sp. HMWF018 (*Betaproteobacteria*), *Burkholderiales* bacterium 68-12 (*Betaproteobacteria*), *Sedimenticola selenatireducens* (*Gammaproteobacteria*), *Ketobacter alkanivorans* (gammaproteobacterial *Oceanospirillales*), and *Aestuarilibacter aggregatus* (*Gammaproteobacteria*), respectively (**Table 2**). In addition, differentially expressed β-oxidation genes were identified in these binned genomes (**Fig. 4A**). Bin13, Bin14, and Bin18 contained genes involved in nitrate and nitrite reduction, chemotaxis, and flagellar synthesis (**Dataset S1**); some of them were differentially expressed **(Table 3).** Differentially expressed hydrolase or esterase genes were not identified in the SPN mesocosms.

**Figure 4.**
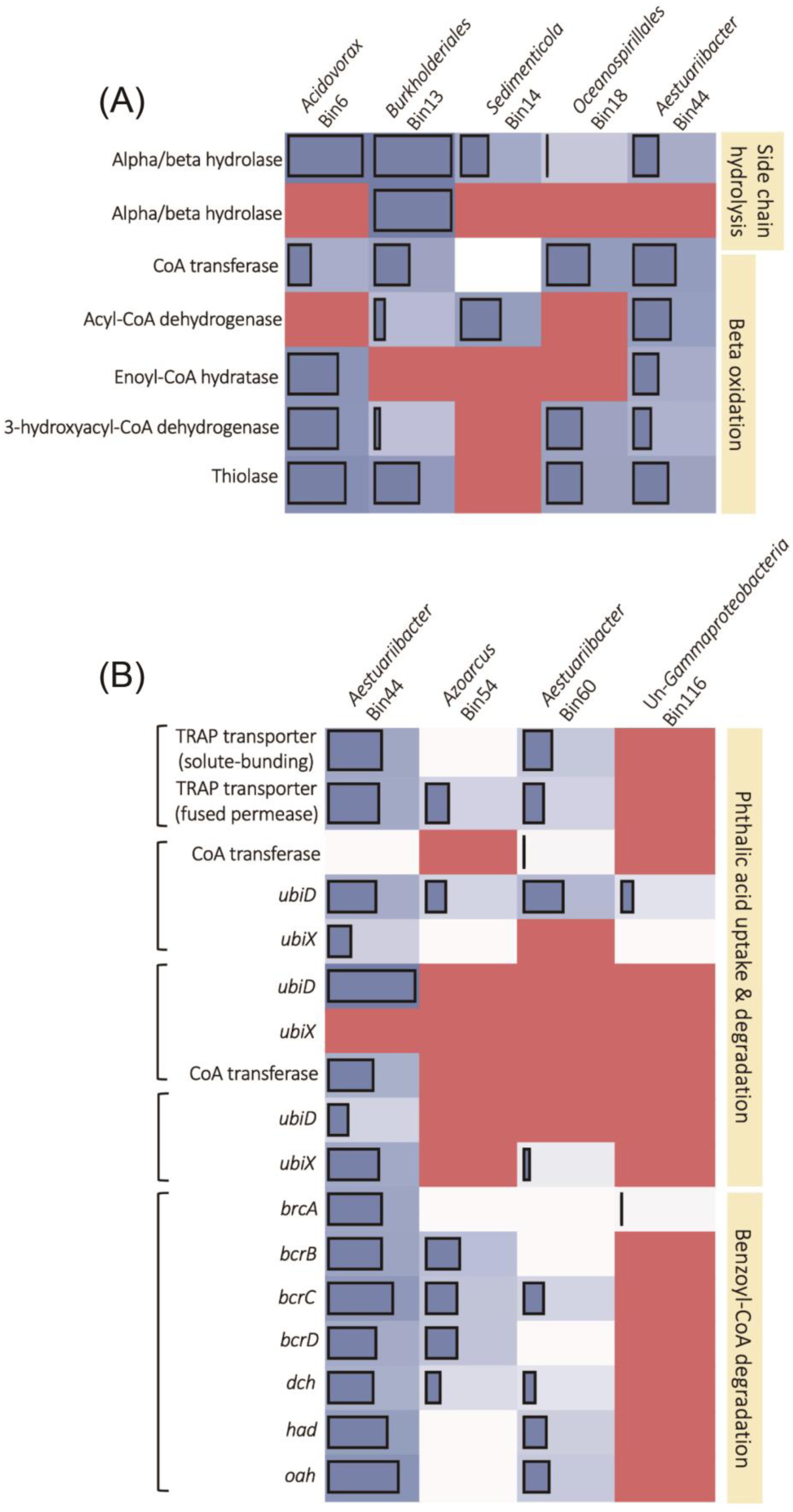
Abundance of differential expression genes involved in **(A)** alkyl side-chain degradation and **(B)** *o***-**phthalic acid uptake and degradation in DEHP-treated sediments. Genes identified in the same contig in each binned genome were grouped in the same square brackets. Blue blocks and bars indicate the differentially expressed genes and their abundance (logCPM). White blocks denote genes that were identified but not differentially expressed. The red block indicates genes not recovered. The contig number, fold change, CPM, and annotation of these genes are listed in **Dataset S1**. Abbreviation: *ubiD*: UbiD-like phthaloyl-CoA decarboxylase, *ubiX*: flavin prenyltransferase, *brcABCD*: benzoyl-CoA reductase, *dch*: dienoyl-CoA hydratase, *had*: hydroxyacyl-CoA dehydrogenase, *oah*: oxoacyl-CoA hydrolase.

**Table 1.** Hydrolase potentially involved in the degradation of DEHP alkyl side-chain in DEHP-treated sediments.

| Gene/accession no. | Genome | InterPro superfamily | Signal peptide |
| --- | --- | --- | --- |
| k141_3815848_2 | *Acidovorax* Bin6 | IPR029058 AB_hydrolase | Secretory signal peptide (1-25) |
| k141_2812817_2 | *Burkholderiales* Bin13 | IPR029058 AB_hydrolase | Secretory signal peptide (1-37) |
| k141_4060503_4 | *Burkholderiales* Bin13 | IPR029058 AB_hydrolase | Secretory signal peptide (1-23) |
| k141_138169_2 | *Sedimenticola* Bin14 | IPR029058 AB_hydrolase | ND |
| k141_2938615_2 | *Ketobacter* Bin18 | IPR029058 AB_hydrolase | ND |
| k141_853245_1 | *Aestuariibacter* Bin44 | IPR029058 AB_hydrolase | lipoprotein signal peptide (1-20) |
| OGB81769.1* | *Burkholderiales bacterium* | IPR029058 AB_hydrolase | Secretory signal peptide (1-23) |
| WP_132980767.1* | *Pigmentiphaga* sp. D-2 | IPR029058 AB_hydrolase | Secretory signal peptide (1-23) |
| WP_130355623.1* | *Pigmentiphaga kullae* | IPR029058 AB_hydrolase | Secretory signal peptide (1-23) |

Parentheses: the position of the signal peptide on the amino acid. ND: not determined
\*Proteins displayed > 70% of amino acid identify to DEHP/MEHP hydrolase (NCB65476) of *Acidovorax* sp. strain 210-6. The details are in the session of phylogenetic analysis

**Table 2.** Features and quality of binned genomes that had partial or complete capacity to degrade DEHP

| Bin | Total length (bp) | # of contigs | N50 | GC (%) | Taxonomy | AAI to closet species (%) | CheckM completeness (%) | CheckM contamination (%) | Degradation capacity |
| --- | --- | --- | --- | --- | --- | --- | --- | --- | --- |
| Bin44 | 1,715,874 | 330 | 10,375 | 48.15 | Gamma-proteobacteria | *Aestuariibacter aggregatus* (84.1) | 46.55 | 26.8 | Alkyl side-chain<br>Phthalic acid |
| Bin6 | 251,465 | 125 | 1,929 | 63.07 | Beta-proteobacteria | *Acidovorax* sp.<br>HMWF018 (97.12) | 4.02 | 0 | Alkyl side-chain |
| Bin13 | 370,293 | 60 | 11,924 | 62.49 | Beta-proteobacteria | *Burkholderiales*<br>bacterium 68-12 (84.61) | 25.86 | 6.43 | Alkyl side-chain |
| Bin14 | 472,849 | 30 | 25,611 | 58.34 | Gamma-proteobacteria | *Sedimenticola selenatireducens* (85.94) | 15.99 | 0 | Alkyl side-chain |
| Bin18 | 843,089 | 125 | 18,585 | 57.57 | Gamma-proteobacteria | *Ketobacter alkanivorans* (65.59) | 24.29 | 6.9 | Alkyl side-chain |
| Bin54 | 1,716,048 | 347 | 8,133 | 64.99 | Beta-proteobacteria | *Azoarcus communis* (74.7) | 66.61 | 26.03 | Phthalic acid |
| Bin60 | 5,123,840 | 411 | 31,029 | 47.30 | Gamma-proteobacteria | *Aestuariibacter aggregatus* (85.9) | 93.65 | 35.95 | Phthalic acid |
| Bin116 | 443,991 | 236 | 1,808 | 56.74 | Gamma-proteobacteria | *Gammaproteobacteria*<br>bacterium BRH_c0 (84.1) | 3.82 | 0.35 | Phthalic acid |

**Table 3.** Differentially expressed genes (KEGG orthologs, KO) related to nitrate reduction, chemotaxis, and motility in the binned genomes.

| Function | *Burkholderiales* Bin13 | *Sedimenticola* Bin14 | *Oceanospirillales* Bin18 | *Aestuariibacter* Bin44 | *Azoarcus* Bin54 | *Aestuariibacter* Bin60 |
| --- | --- | --- | --- | --- | --- | --- |
| Nitrate reduction | nitrate reductase/nitrite oxidoreductase, alpha & beta subunit (K03070, K03071)<br>nitrate reductase molybdenum cofactor assembly chaperone (K00373)<br>nitrate reductase gamma subunit (K00374) | nitrate/nitrite transporter (K02575 & K02576)<br>nitrate reductase/nitrite oxidoreductase, beta subunit (K03071) | periplasmic nitrate reductase NapA (K2567) | nitrite reductase large subunit (K00362)<br>nitrate reductase/nitrite oxidoreductase, alpha & beta subunit (K03070, K03071)<br>nitrate reductase catalytic subunit (K00372)<br>nitrate reductase molybdenum cofactor assembly chaperone (K00373)<br>nitrate reductase gamma subunit (K00374)<br>nitrate/nitrite transporter (K02575) | periplasmic nitrate reductase NapA (K2567)<br>nitrate reductase/nitrite oxidoreductase, beta subunit (K03071)<br>nitrate reductase molybdenum cofactor assembly chaperone (K00373) | nitrate reductase/nitrite oxidoreductase, alpha & beta subunit (K03070, K03071)<br>nitrate reductase molybdenum cofactor assembly chaperone (K00373)<br>nitrate reductase gamma subunit (K00374)<br>nitrate/nitrite transporter (K02575) |
| Chemotaxis | Not recovered | Recovered but not differentially expressed | chemotaxis methyltransferase CheR (K00575)<br>Chemotaxis glutaminase (K2390)<br>methyl-accepting chemotaxis protein (K03406)<br>chemotaxis sensor kinase CheA (K03407)<br>chemotaxis protein CheVY (K03415, K03413)<br>chemotaxis pili protein ChpABC (K06596, K06597, K06598) | Not recovered | methyl-accepting chemotaxis protein (K03406)<br>chemotaxis sensor kinase CheA (K03407)<br>Chemotaxis glutaminase (K2390)<br>Chemotaxis protein CheWYV (K03408, K03413, K03415) | chemotaxis methyltransferase CheR (K00575)<br>Chemotaxis glutaminase (K2390)<br>chemotaxis protein MotB (K02557)<br>methyl-accepting chemotaxis protein (K03406)<br>chemotaxis sensor kinase CheA (K03407)<br>purine-binding protein CheW (K3408)<br>chemotaxis protein CheD (K3411) |
| Motility | flagellin (K02406) | Recovered but not differentially expressed | hook-associated protein 123 (K02396, K02407, K02397)<br>biosynthesis protein FlgF (K02404)<br>flagellin (K02406)<br>flagellar protein FliL (K02415)<br>biosynthesis protein FlhG (K04562) | flagellin (K02406)<br>hook-associated protein 2 (K02407)<br>flagellar protein FlaG (K06603)<br>flagellar protein FliS (K02422) | basal-body rod protein FlgBD (K02387, K02389)<br>Biosynthesis protein FlgN (K02399)<br>flagellin (K02406)<br>hook-associated protein 2 (K02407)<br>flagellar protein FlaG (K06603)<br>flagellar brake protein (K21087) | biosynthesis protein FlhAF (K02400, K02404)<br>M-ring protein FliF (K02409)<br>Flagellum ATP synthase (K02412)<br>hook-length control FliK (K02414)<br>fmotor switch protein FliMN (K02416, K02417)<br>biosynthesis protein FlhG (K04562) |

### Binned genomes with complete *o*-phthalic acid degradation capacity in the DEHP-treated mesocosms

In the SDN mesocosms, 23 genes belonging to the UbiD-like decarboxylase family were upregulated. According to blastp results, only eight UbiD homologs identified in eight contigs share high amino acid sequence identity (AAI; 74% to 85%) to phthaloyl-CoA decarboxylase of *A*. *aromaticum* EbN1=DSM19081, *A*. *toluclasticus* ATCC 700605, and *T*. *chlorobenzoica* 3CB-1=DSM18012 (**Dataset S1**). Furthermore, we discovered genes annotated as flavin prenyltransferase (UbiX) [22], CoA-transferase, TRAP transporter, and transposase in some of these contigs (**Dataset S1**); however, none of these genes were differentially expressed in the SDN mesocosms.

We selected binned genomes containing the phthaloyl-CoA decarboxylase gene to determine whether they could anaerobically degrade *o*-phthalic acid. Up to 394 binned genomes were generated by MaxBin 2.0, but 115,543 contigs were un-binned. We observed that three contigs with phthaloyl-CoA decarboxylase genes were binned into Bin44, whereas in other binned genomes—Bin54, Bin60, and Bin116—only a single contig with the phthaloyl-CoA decarboxylase gene was binned (**Fig. 4B**). However, two contigs—K141_1830739 and K141_260687—with this gene could not be binned (**Dataset S1**).

Two genes involved in *o*-phthalic acid uptake were also identified in these binned genomes (except for Bin116) and two un-binned contigs. Moreover, crucial genetic components for anaerobic benzoyl-CoA degradation—including benzoyl-CoA reductase (*brcABCD*), dienoyl*-*CoA hydratase (*dch*), hydroxyacyl-CoA dehydrogenase (*had*), and oxoacyl-CoA hydrolase (*oah*)—were identified in Bin44, Bin54, and Bin60; in Bin116, only one gene encoding benzoyl-CoA reductase subunit A (*brcA*) was recovered (**Fig. 4B** and **Dataset S1**). In addition, we noted that most of the genes related to nitrate reduction, chemotaxis, and flagellar synthesis were differentially expressed in Bin44, Bin54, and Bin60 (**Table 3** and **Dataset S1**).

The closest taxonomic affiliation of Bin44, Bin54, Bin60, and Bin116 was *Aestuariibacter aggregatus* (*Gammaproteobacteria*), *Azoarcus communis* (*Betaproteobacteria*), *Aestuariibacter aggregatus*, and unclassified *Gammaproteobacteria bacterium* BRH_c0, respectively. The CheckM result revealed that the Bin60 and Bin116 genomes showed the highest (93.65%) and lowest degrees of completeness, respectively. The overall information and quality of these selected binned genomes are detailed in **Table 2**. We also recovered *Thauera* Bin 97 and *Azoarcus* Bin394 with transport and degradation capacity for *o*-phthalic acid in the *o*-phathalic acid-treated sediments (**Table S2**). The details are described in supplementary information.

### *Acidovorax* sp. strain 210-6 isolated from the DEHP-enriched mesocosm displayed degradation capacity for the alkyl side-chains of DEHP

To elucidate the functionality of the DEHP-enriched sediment bacteria, we cultured and isolated the DEHP degrader *Acidovorax* sp. strain 210-6 from the SDN1 mesocosm and sequenced its genome (accession no.: GCA_010020825.1). The growth curve measurement and metabolite profile analysis showed that strain 210-6 could utilize DEHP, MEHP, and 2-ethyl-1-hexanol as sole carbon and energy sources in a denitrifying medium (**Fig. 5**). Notably, we observed the accumulation of *o*-phthalic acid in the DEHP- and MEHP-fed cultures concomitant with the consumption of these substrates (**Fig. 5A**). The OD at 600 nm did not increase when either *o-*phthalic acid or benzoic acid was present as the sole carbon and energy sources, and these substrates were not apparently consumed.

**Figure 5.**
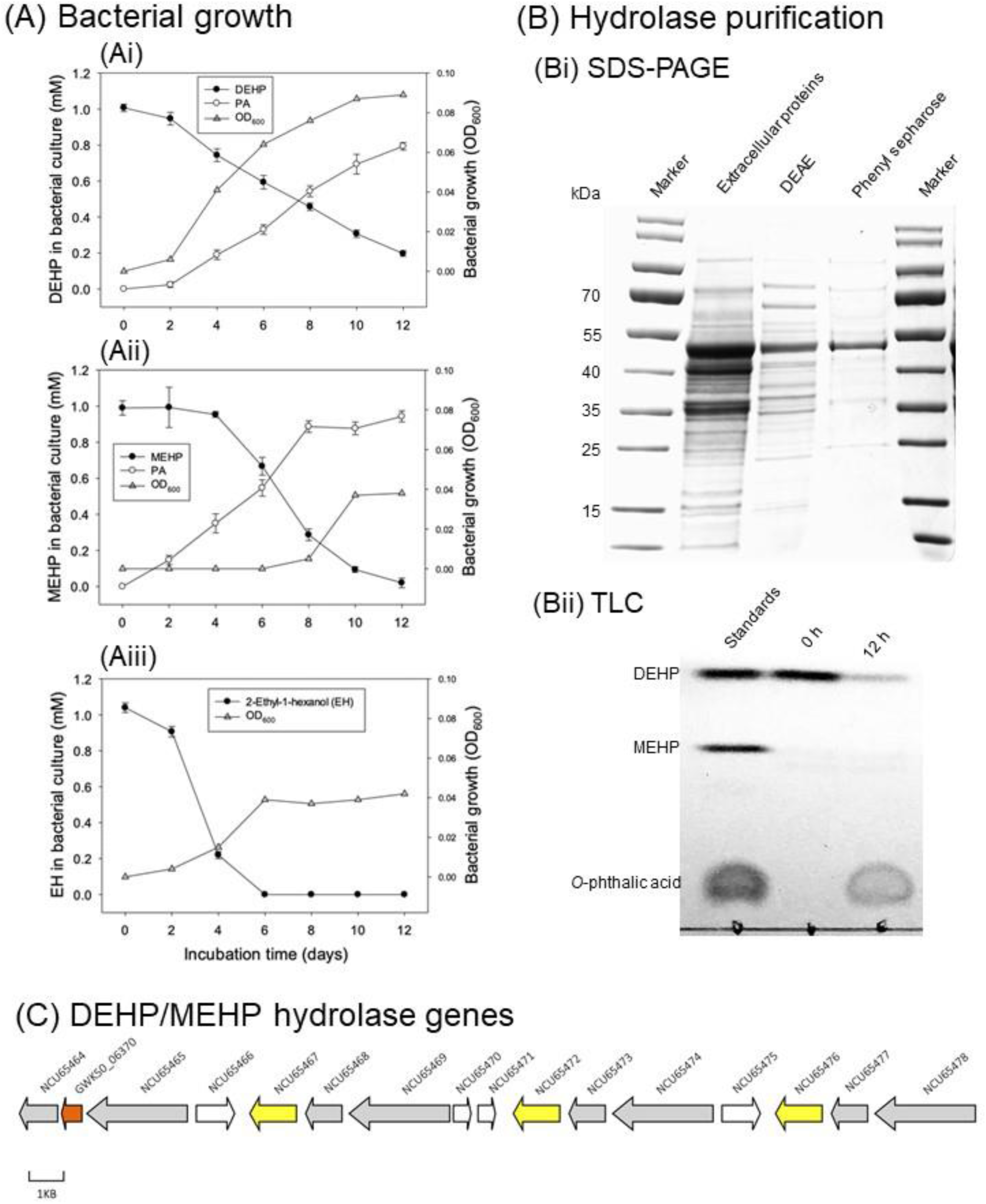
Functional characterization of *Acidovorax* sp. strain 210-6 and its DEHP/MEHP hydrolase. (A) Anaerobic growth of the strain 210-6 with DEHP (Ai), MEHP (Aii), and EH (Aiii). The DEHP, MEHP, and *o*-phthalic acid (PA) were quantified using UPLC–APCI– HRMS, whereas EH was quantified using GC–MS. (B) Purification and characterization of DEHP/MEHP hydrolase. (Bi) SDS-PAGE (4%–12%) indicated the purification of the DEHP/MEHP hydrolase from the extracellular proteins of strain 210-6. (Bii) TLC indicated the hydrolase activity of the purified protein toward DEHP. The detected major product was PA but not MEHP. After 12 h of anaerobic incubation, metabolites were extracted with ethyl acetate, separated through TLC, and visualized by spraying the TLC plate with 30% (v/v) H_2_SO_4_. (C). Presence of three identical copies of hydrolase genes (NCU65467, NCU65472, and NCU65476) in the chromosome of strain 210-6. Protein ID (NCU_) or locus tag (GW50_) are present on each gene. Yellow: DEHP/MEHP hydrolase. Grey: transposase/integrase. White: hypothetical proteins. Red: proteins not related to DEHP degradation. GWK50_06370: FAD-dependent oxidoreductase.

The strain 210-6 genome contained three identical copies of 16S rRNA genes, which displayed 97.7% sequence similarity to the 16S rRNA gene of *Acidovorax valerianellae* DSM 16619. The chromosome and plasmid harbor 29 genes encoding alpha/beta hydrolase. Consistent with their growth profiles, genes involved in the degradation of *o-*phthalic acid were not identified in the genome of strain 210-6. We did not observe any DEHP degradation activity when we used the soluble proteins of *Acidovorax* sp. 210-6. However, SDS-PAGE analysis of the active fraction obtained from extracellular proteins showed a protein band corresponding to an approximate molecular mass of 50 kDa (**Fig. 5BI**). The purified protein could transform DEHP into *o*-phthalic acid within 12 h, underscoring its side-chains hydrolysis activity. Moreover, the temporal accumulation of MEHP was not observed, suggesting the hydrolysis of DEHP into MEHP as the rate-limiting step. The results of liquid chromatography (LC)–tandem mass spectrometry (MS/MS) analysis revealed that one alpha/beta hydrolase, encoded by the chromosomal gene NCU65476, displayed the highest posterior error probabilities (PEP) score for tryptic peptides originating from the active fraction. Notably, NCU65476 and its two identical copies, namely NCU65467 and NCU65472, were located in the same gene cluster on the chromosome (**Fig. 5C**); each gene carried predicted signal peptide and was flanked by transposase elements.

### Phylogenetic analysis of phthaloyl-CoA decarboxylases and DEHP/MEHP hydrolases

The unrooted maximum likelihood tree of phthaloyl-CoA decarboxylase showed that 8 genes annotated as phthaloyl-CoA decarboxylase in the gammaproteobacterial bins (except for k141_3428055_1, k141_2680687_1, and k141_1830739_7) formed a distinct clade, whereas another phthaloyl-CoA decarboxylase gene (k141_1104744_1) recovered in *Azoracus* Bin394 was placed in the same clade with phthaloyl-CoA decarboxylase genes derived from denitrifying *Betaproteobacteria* (*A*. *aromaticum* DSM19081*, A*. *toluclasticus* ATCC 700605, *T*. *chlorobenzoica* DSM18012, and *Azoarcus* sp. PA01). These two clades were distinct from the phthaloyl-CoA decarboxylase of sulfate-reducing *Deltaproteobacteria* (*Desulfosarcina cetonia* DSM 7267, and *Desulfobacula toluolica* DSM7467; **Fig. 6**). Other UbiD family decarboxylases—namely 2,5-furandicarboxylate decarboxylase, 3-octaprenyl-4-hydroxybenzoate decarboxylase, phenolic acid decarboxylase subunit C, phenylphosphate carboxylase subunit alpha, and phenylphosphate carboxylase subunit beta—were placed into the same clades along with their homologs identified in DEHP-treated mesocosms (**Fig. S5**). In addition, we noted that UbiD from *Acidovorax* sp. strain 210-6 (protein id: NCU67919) was grouped into the clade with other 3-octaprenyl-4-hydroxybenzoate decarboxylase sequences in line with its inability to degrade *o*-phthalic acid.

**Figure 6.**
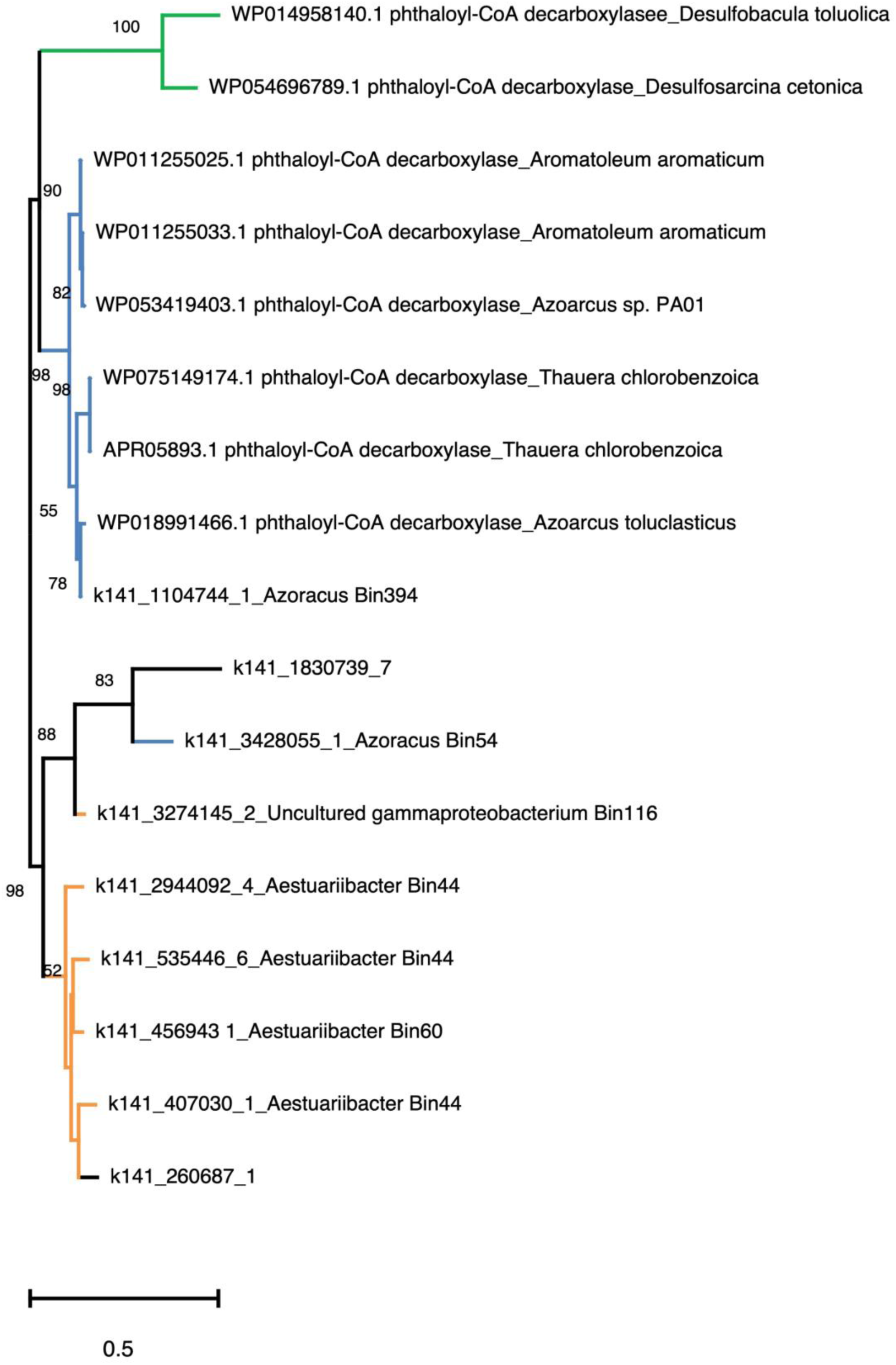
Maximum likelihood tree of phthaloyl-CoA decarboxylase from bacterial isolates and in DEHP-treated sediments. Orange branch: gammaproteobacterial nitrate-reducing phthalic acid degraders. Blue branch: betaproteobacterial nitrate-reducing *o*-phthalic acid degraders. Green branch: deltaproteobacterial sulfate-reducing phthalic acid degraders. Branch support of higher than 50% of the bootstrapping time is shown.

In the unrooted maximum likelihood tree of hydrolases (**Fig. 7**), hydrolases involved in the aerobic side-chain degradation of monoalky PAEs (from aerobic *Rhodococcus* spp. and *Gordonia* spp.) formed a distinct lineage. Most of the hydrolases involved in the degradation of dialkyl PAEs from several aerobes—*Acinetobacter* sp. M673, *Gordonia* sp. YC-JH1, *Sulfobacillus Acidophilus* DSM1032, and *Sphingobium* sp. SM42 and PAMC26605—were also placed into the same clade, except for hydrolases involved in side-chains degradation of di-*n*-butyl phthalate (from *Sphingobium* sp. SM42) and DEHP (from *Gordina* sp. 5F). By contrast, we observed that all DEHP/MEHP hydrolases from denitrifying bacteria and bins formed a distinct clade. Moreover, four DEHP/MEHP hydrolases (NCU68007, NCU65467, NCU65472, and NCU65476) from *Acidovorax* sp. strain 210-6, three putative hydrolases (K141_4060503_4, K141_3815848_2, and K141_2812817_2) from betaproteobacterial bins, as well as three hypothetical proteins from the two strains of *Pigmentiphaga* and uncultured *Burkholderiales* bacterium were in the same lineage. These hypothetical proteins also belonged to the alpha/beta hydrolase family IPR029058 with predicted signal peptides **(Table 1).** Other putative DEHP hydrolases—K141_138169_2, K141_2938615_2, and K141_853245_1—derived from gammaproteobacterial bins were in different lineage.

**Figure 7.**
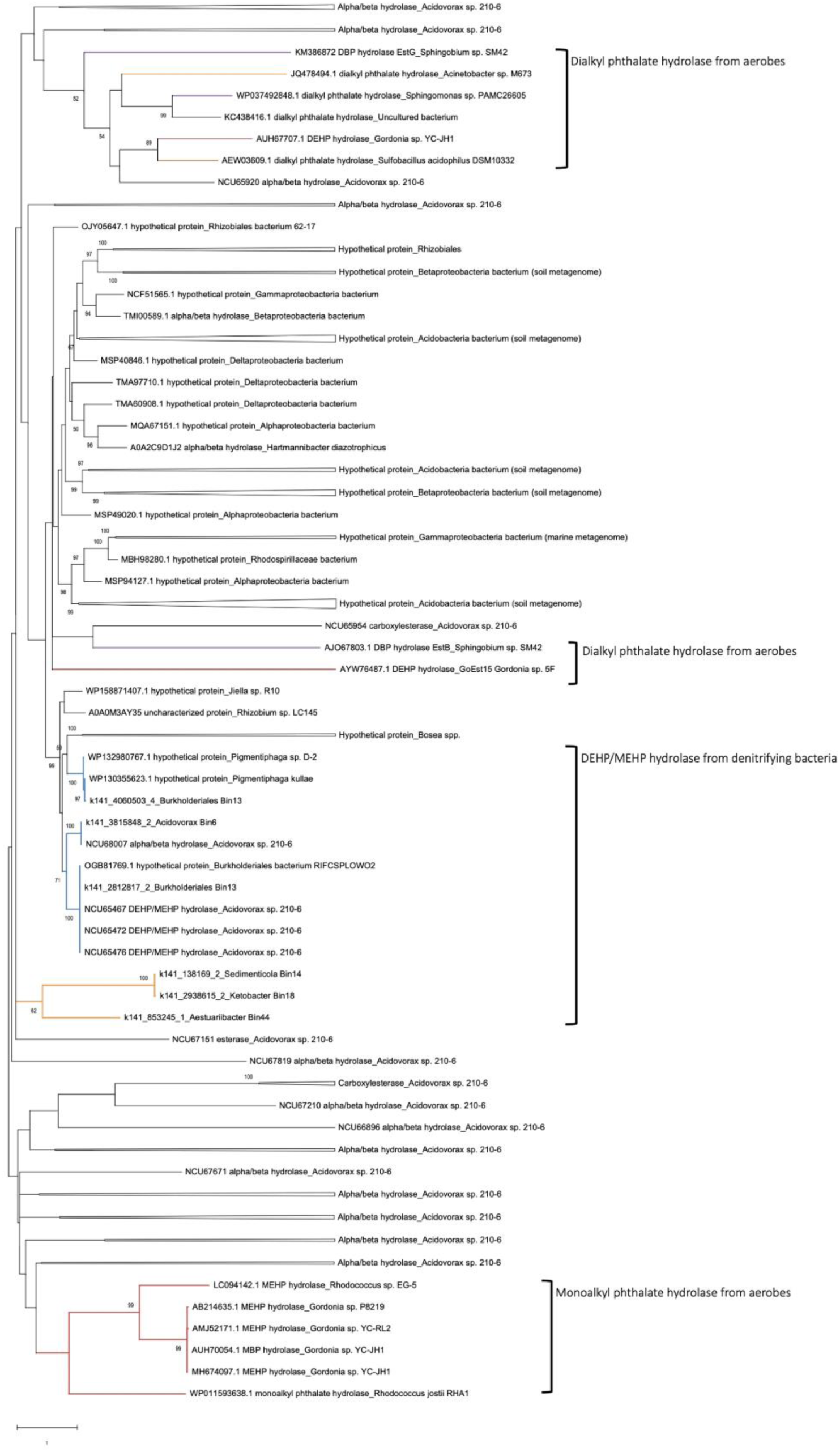
Maximum likelihood tree of hydrolases involved in degradation of alkyl side-chains on DEHP and other PAEs and other hydrolases from *Acidovorax* sp. strain 210-6. The phylogeny of the protein sequences from the NCBI and uniport with an identify ≥ 40% to the NCU65467 of *Acidovorax* sp. strain 210-6 was also inferred. Purple branch: aerobic alphaproteobacterial degraders. Blue branch: anaerobic betaproteobacterial degraders. Red branch: aerobic actinobacterial degraders. Orange branch: anaerobic gammaproteobacterial degraders. Brown branch: aerobic Firmicutes degraders. Branch support of more than 50% of the bootstrapping time is shown.

### Transposon genes in contigs and gene clusters with key enzymes for DEHP degradation

In *Burkholderiales* Bin13, we noted that two putative DEHP/MEHP hydrolase genes were flanked by transposon genes in two contigs (**Fig. 8**). Moreover, hydrolase genes in the genomes of the *Burholderiales bacterium* RIFCSPLOWO2 and *Pigmentiphapa* sp. D-2 that displayed high AAI (99.7 % and 72.2 %, respectively) to the DEHP/MEHP hydrolase gene (NCU65476) of *Acidovorax* sp. strain 210-6 were flanked by transposon genes (**Fig. 8**). A similar arrangement was observed in *Aestuariibacter* Bin44, in which genes involved in anaerobic *o*-phthalic acid degradation—namely flavin prenyltransferase, phthaloyl-CoA decarboxylase, and CoA transferase—were adjunct to transposon elements (**Fig. 8**)

**Figure 8.**
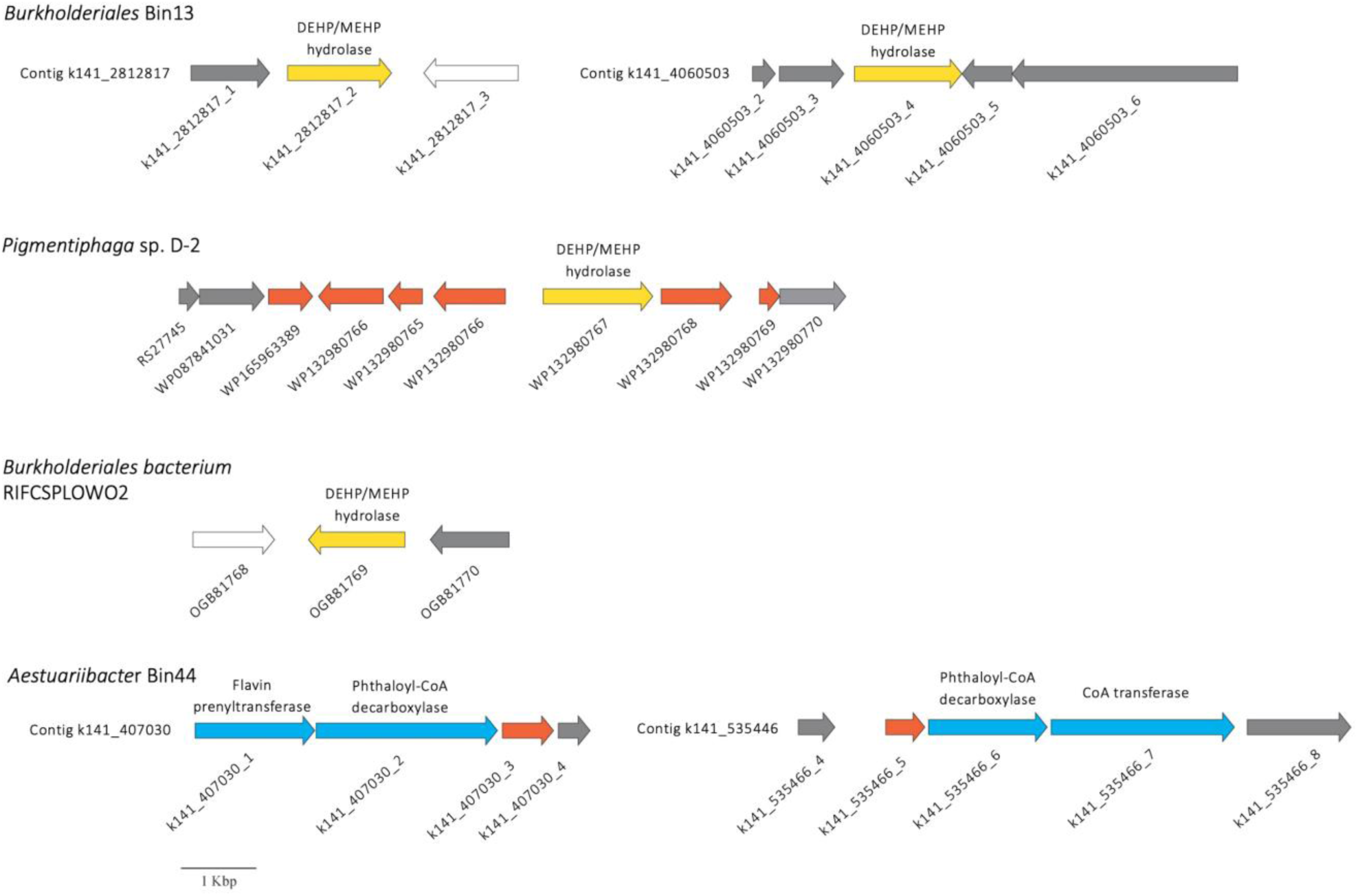
Transposase genes in contigs or gene clusters carrying essential components for side-chain hydrolysis of DEHP and *o*-phthalic acid degradation. Grey: transposase/integrase. Yellow: DEHP/MEHP hydrolase. Blue: genes for *o*-phthalic acid degradation. White: hypothetical proteins. Red: proteins not related to DEHP degradation. Sequence references of genomes of *Pigmentiphaga* sp. D-2 and *Burkholderiales* bacterium RIFCSPLOWO2 are indicated. Full annotation of Bin13 and Bin44 is listed Dataset S1.

## Discussion

In this study, we used an integrated multi-omics approach to identify DEHP degraders and elucidate the degradation mechanisms in urban estuarine sediments containing much accumulated DEHP. Several lines of evidence suggest that DEHP degradation in estuarine sediments mainly occurs through a synergistic microbial metabolism: (i) in DEHP-treated mesocosms, various betaproteobacteria and gammaproteobacteria were enriched, and (ii) most of the binned genomes lacked the complete set of degradation genes for DEHP; instead, these bins possessed genes either for alkyl side-chains or for *o*-phthalic acid degradation. A similar observation was made in sediment isolate *Acidovorax* sp. strain 210-6, which could degrade only alkyl side-chains but not *o-*phthalic acid. This finding is different from those of most biodegradation studies, which have indicated that the uptake and mineralization of organic micro-pollutants are carried out by single bacterial isolates. For example, bacterial community structure analysis indicates that the complete mineralization of endocrine-disrupting steroids in sludge and sediment mesocosms is achieved by a single betaproteobacterial genus [23–25]. Moreover, Moreover, several denitrifying bacteria capable of complete steroid degradation have been isolated from steroid-treated mesocosms [26].

Synergistic networks between diverse bacteria are reportedly required for complete DEHP degradation [27–29]; however, the corresponding studies have not fully elucidated the biochemical mechanisms involved in this bioprocess or provided insights into the degradation role of each microorganism. In our DEHP-treated mesocosms, six putative DEHP hydrolase genes (four with a predicted signal peptide) were differentially expressed, and one of the hydrolases, k141_3815854_2, was identified in Bin6, which displayed the closest AAI to *Acidovorax* sp. HMWF018. This hydrolase exhibited 100% AAI to a putative DEHP/MEHP hydrolase gene (NCU68007; **Dataset S1**) on the plasmid of strain 210-6, suggesting that *Acidovorax* Bin6 is a DEHP side-chains degrader. Furthermore, the isolation and characterization of strain 210-6 confirmed *Acidovorax* spp. as active DEHP degraders in the estuarine sediments. Putative DEHP-degrading hydrolase (k141_138169_2) and β-oxidation genes for anaerobic 2-ethylhexanol degradation were identified in Bin14, and its closest taxonomy affiliation was *Sedimencola selenatireducens*. The degradation of DEHP and other PAEs has not been previously reported for the genus *Sedimenticola* [30–33]. Their apparent enrichment in DEHP-treated sediments suggests that *Sedimenticola* plays a role in DEHP degradation. The key genes involved in anaerobic *o*-phthalic acid degradation were not recovered in *Sedimencola* Bin14; thus, we speculated that *Sedimenticola* may also be involved in the alkyl side-chains degradation of DEHP in denitrifying sediments.

On the basis of the profiles of DEHP-derived metabolites, changes in community structures, as well as the taxonomy and functionality of binned genomes, we propose that in the examined denitrifying sediments, complete DEHP degradation requires synergistic metabolism. *Acidovorax* (Bin6), unclassified betaproteobacterium *Burkholderiaceae* (Bin13), *Sedimenticola* (Bin14), and *Oceanospirillales* member SS1_B_06_26 (Bin18) are major degraders of the alkyl side-chains (namely the 2-ethylhexanol moiety) of DEHP. The remaining *o*-phthalic acid is likely transported into bacterial cells by a specific TRAP transporter and primarily degraded by uncultured gammaproteobacterial MBAE14 (Bin116) and *Aestuariibacter* (Bin60) through the anaerobic benzoyl-CoA pathway (**Fig. 9**). Synergistic interactions in a bisphenol A-degrading microbial community were also identified in a recent study [34], suggesting the crucial role of synergistic microbial metabolism in removing endocrine-disrupting aromatics from contaminated environments.

**Figure 9.**
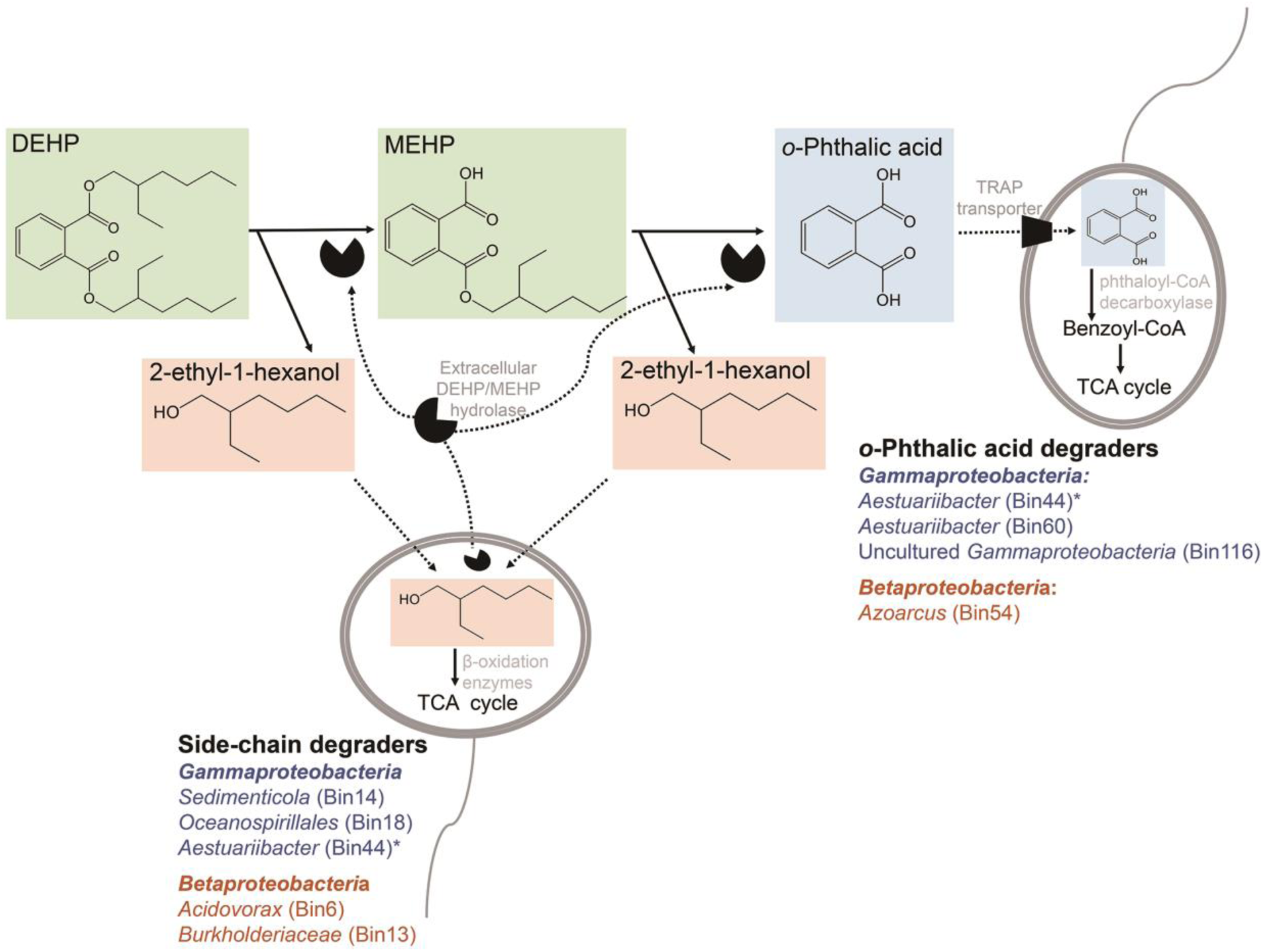
Diagram of proposed DEHP degradation through microbial synergistic metabolism in denitrifying sediments. The betaproteobacterium *Acidovorax* and gammaproteobacterium *Sedimenticola* are major degraders of DEHP side-chains. The resulting *o*-phthalic acid is then imported through a specific TRAP transporter and degraded by other betaproteobacteria (e.g., *Azoarcus*) and gammaproteobacteria (e.g., *Aestuariibacter*). *The gammaproteobacterial Bin44 has degradation capacity for both side-chains and *o*-phthalic acid. Orange: *Gammaproteobacteria*; Blue: *Betaproteobacteria*.

Our community and metagenomic analyses suggest that *Thauera* and *Azoarcus* are ecologically relevant degraders for *o*-phthalic acid [17,19,20]. Unexpectedly, our phylogenetic and metagenomic analyses indicated that *o*-phthalic acid degraders in DEHP-treated sediments are mainly denitrifying gammaproteobacteria, revealing that the diversity of anaerobic *o*-phthalic acid degraders is not only restricted to denitrifying betaproteobacteria and sulfate-reducing deltaproteobacteria such as *Desulfosarcina cetonia* DSM 7267 and *Desulfobacula toluolica* Tol2=DSM7467 [35]. Moreover, whether DEHP is toxic to *Thauera* and *Azoarcus* remains unclear. An apparent decrease in the abundance of other bacteria (e.g., *Clostridiales*) can be observed due to their sensitivity to DEHP [36–38]. In addition, differentially expressed genes related to chemotaxis and flagellar synthesis in DEHP-degrading *Sedimenticola* (Bin 14) and *Aestuariibacter* (Bin44) as well as *o*-phthalic acid-degrading *Aestuariibacter* (Bin60) suggest that DEHP can be an attractant for not only DEHP degraders but also some *o*-phthalic acid-degrading anaerobes. Some bacteria display chemotaxis toward substrates that they are not able to degrade [39, 40]. Accordingly, we assumed that DEHP induced chemosensory in some *o*-phthalic acid-degrading gammaproteobacteria, enhancing their better attraction to *o*-phthalic acid produced from DEHP.

Chemotaxis has been proposed to facilitate horizontal gene transfer because the ability of bacteria to accumulate around pollutant substrates increases the possibility of the transfer of relevant catabolic genes [41]. Transposon elements have been identified in the gene clusters involved in DEHP side-chains hydrolysis in the genome of *Acidovorax* sp. strain 210-6 as well as in *o*-phthalic acid degradation in *A*. *aromaticum* DSM19081 (on plasmid), *Azoarcus* sp. PA01 (on chromosome), and *T*. *chlorobenzoate* DSM18012 (on chromosome) [19]. Furthermore, Sanz [42] suggested that the complete *o*-phthalic acid degradation pathway in denitrifying bacteria, together with the *o*-phthalic acid transporter gene, is successfully transferred to heterologous bacteria that are unable to use *o*-phthalic acid as a substrate. In the present study, a similar observation was made in DEHP side-chains-degrading *Burkholderiales* Bin13 and *o*-phthalic acid-degrading *Aestuariibacter* Bin44. Surprisingly, the gene cluster or contig with potential DEHP-degrading hydrolases from *Pigmentiphaga* sp. D-2 and uncultured *Burkholderiales* bacterium also contained a transposase gene. These findings indicate that in the natural environment, DEHP degradation capacity may be widespread among proteobacteria due to transposon-mediated horizontal gene transfer [43]; this further supports the notion that anthropogenic inputs may not only result in changes in community structures but also in ecosystem functioning [44, 45].

## Materials and methods

### Sample site and sediment collection

In the northern part of Taiwan, up to six million people reside in the Taipei City located in the basin of the Tamsui River and Keelung River. Our sampling site, Guandu estuary (25°6’59.56″N, 121°27’46.99″E) is located downstream of these two rivers and receives sewage discharges and waste effluent from the Taipei metropolitan area. The profile of PAEs in river ecosystems in Taiwan has been identified; the average DEHP concentration in Tamsui River sediments was 2.3 µg/g; other PAEs–namely diethyl phthalate (DEP), dipropyl phthalate (DPP), di-*n*-butyl phthalate (DBP), diphenyl phthalate (DPhP), benzylbutyl phthalate (BBP), dihexyl phthalate (DHP), and dicyclohexyl phthalate (DCP)–were either not detected or were detected in concentrations lower than 0.5 µg/g [46]. The subsurface sediments (5–10 cm) of Guandu estuary were abundant in nitrate and denitrifying bacteria [47]. During the low tide that occurred on July 2, 2018, two independent subsurface sediment and river water samples were collected as described previously [24]. These samples were stored at 4 °C and transported to the laboratory within 1 hour for performing mesocosm incubation of denitrifying sediments.

### Mesocosm incubation of denitrifying sediments with DEHP, benzoic acid or *o*-phthalic acid

Mesocosm experiments were performed in 1-L sterilized serum bottles containing subsurface sediments (approximately 200 g) and river water (approximately 800 mL). Four types of mesocosm (two replicates in each)–10mM of sodium nitrate and 1mM of DEHP (SDN mesocosm), 10mM of sodium nitrate and 1mM of *o*-phthalic acid (SPN mesocosm), 10mM of sodium nitrate and 1mM of benzoic acid (SBN mesocosms), and 10mM of nitrate only (SN mesocosm as the control)–were prepared under the anoxic condition by purging 80 % (vol/vol) of nitrogen gas and 20% (vol/vol) of carbon dioxide into serum bottles sealed with butyl rubber stoppers. All mesocosms were further reduced with 0.5 mM Na_2_S to neutralize oxygen and incubated at 25°C with agitation. The consumption of nitrate was monitored using the Spectroquant nitrate test kit HC707906 (Merck, Germany). Sodium nitrate (10 mM) was resupplied when nitrate was depleted until the exogenous substrates were completely degraded. Samples of river water and sediment mixture (approximately 10 mL) were obtained from each mesocosm every 2 or 3 days, stored at −80°C for metabolite extraction, and preserved in LifeGuard Soil Preservation Solution (Qiagen, Germany) for total DNA/RNA extraction. All chemicals were purchased from Sigma-Aldrich, Merck KGaA (St. Louis, Missouri, USA).

### UPLC–APCI–HRMS analysis of DEHP-derived metabolites

Hydrophobic non-CoA thioester metabolites in the mesocosms (SDN, SPN, and SBN samples) were extracted using ethyl acetate as described previously [24]. Crude extracts were then applied in UPLC**–**MS with UPLC coupled to an APCI**–**mass spectrometer to identify and quantify metabolites. DEHP and its metabolites were first separated using a reversed-phase C_18_ column (Acquity UPLC® BEH C18; 1.7 μm; 100 × 2.1 mm; Waters) at a flow rate of 0.3 mL/min at 65°C (oven temperature). The mobile phase was a mixture of two solutions: solution A [0.1% formic acid (v/v) in 2% acetonitrile] and solution B [0.1% formic acid (v/v) in isopropanol]. Separation was achieved using a gradient of solvent B from 1% to 99% over 7 min. APCI–MS analysis was performed using an Orbitrap Elite Hybrid Ion Trap-Orbitrap Mass Spectrometer (Thermo Fisher Scientific, Waltham, MA, USA) equipped with a standard APCI source. MS data were collected in the positive ionization mode (parent scan range: 100–500 *m/z*). The capillary and APCI vaporizer temperatures were 120°C and 400°C, respectively; the sheath, auxiliary, and sweep gas flow rates were 40, 5, and 2 arbitrary units, respectively. The source voltage was 6 kV, and the current was 15 μA. The elemental composition of individual adduct ions was predicted using Xcalibur™ Software V2.2 (Thermo Fisher Scientific). The following authentic standards were purchased from Sigma-Aldrich: DEHP, MEHP, *o*-phthalic acid and benzoic acid.

### Gas chromatography (GC)–MS analysis of 2-ethyl-1-hexanol

Quantification of the remaining 2-ethyl-1-hexanol in bacterial cultures was performed through GC on an HP 5890 Series II GC device coupled to 5972 Series Mass-Selective Detector (Hewlett-Packard, Palo Alto, CA, USA). A fused-silica capillary GC column (DB-1ms; 60 m × 0.25 mm i.d.; Agilent J &W Scientific, Folsom, USA) chemically bonded with a 100% dimethylpolysiloxane stationary phase (0.25-mm film thickness) was used. The sample was inserted in the spitless mode by using helium as a carrier gas. Initially, the oven temperature was maintained at 60°C and then increased to 95°C at the rate of 2°C/min. Once the temperature had been maintained at 95°C for 1 min, it was increased to 120°C at a rate of 3 °C/min and then maintained at 120°C for 2 min.

Subsequently, the temperature was further increased to 180°C at the rate of 6°C/min and maintained at 180°C for 2 min. Electron impact ionization was used as an ionization source for the GC–MS analysis at 70 eV. Data acquisition was performed in full scan mode from 50 to 300 *m*/*z* over a scan duration of 0.5 s. *n*-Hexane (as blank) was run between samples to remove contamination. Mass calibration was performed using perfluorotributylamine. Authentic standard 2-ethyl-1-hexanol was purchased from Sigma-Aldrich.

### DNA extraction and 16S rRNA gene amplicon sequencing

The bacterial community structure of each mesocosm and temporal changes (day 0 to day 25 for the SN, SBN, and SPN mesocosms and day 0 to day 30 for the SDN mesocosm; two replicates for each) were identified using an Illumina Miseq platform. DNA was extracted from treated sediments by using the PowerSoil DNA isolation kit (Qiagen, Germany). The 16S amplicon libraries targeting the V3-V4 regions of 16S rRNA genes [48] were prepared according to the Illumina 16S metagenomic sequencing library preparation guide by using a gel purification approach as described previously [23]. In total, 96 libraries were generated and their profiles were analyzed using the BioAnalyzer 2100 with a High-Sensitivity DNA kit (Agilent, USA). To ensure the evenness of library pooling, all libraries were subjected to quantitative PCR (qPCR) for normalization by using a Kapa library quantification kit to obtain molar concentrations. For sequencing, the pooled library was run on an Illumina MiSeq sequencer with MiSeq reagent kit V3 (paired end; 2 by 300 bp).

### Bioinformatics processing and taxonomic assignment for 16S amplicon sequencing

MiSeq sequencing generated 27,848,006 reads from 96 sediment samples. USEARCH v11 [49] was used for paired reads assembly, quality filtering, length trimming, and UPARSE OTU clustering [50]. The representative sequences of OTUs were taxonomically assigned against Silva release 132 [51] by using mothur v1.41.3 [52]. Silva release 132 re-classifies *Betaproteobacteria* as an order of *Gammaproteobacteria*. For readability, the abundance of *Gammaproteobacteria* and *Betaproteobacteria* was calculated separately. Less abundant OTUs from all samples (count <2 per OTU with a prevalence of <20% in 96 samples) were removed and then normalized through cumulative sum scaling [53] by using MicrobiomeAnalyst [54]. Similarities between microbial communities among differently treated sediments (SN, SBN, SPN, and SDN) were determined by performing principal-coordinate analysis (PcoA) in MicrobiomeAnalyst on the basis of the weighted UniFrac distance matrix (genus level). The UniFrac distance was calculated according to the neighbor-joining phylogenetic tree file generated by the MUSCLE algorithm [55]. To determine the degraders for benzoic acid, *o*-phthalic acid, and DEHP, we first selected a large increase in relative abundance at bacterial class level. We applied the analysis of variance test ANOVA in MetaboAnalyst 4.0 [56] to identify the bacterial genera in selected classes exhibiting the highest differences in abundance among different mesocosms.

### Metagenomic and metatranscriptomic sequencing

To identify genes responding to exogenous DEHP and *o*-phthalic acid in denitrifying sediments, the total DNA and RNA of SDN1_day14, SDN2_day14, SPN1_day7, and SPN2_day7 were used for metagenome and metatranscriptome analyses. Total DNA and RNA were extracted using the PowerSoil DNA isolation kit and Rneasy PowerSoil total RNA kit (Qiagen, Germany), respectively. For shotgun metagenomic sequencing, the quality of DNA was examined using the Fragment Analyzer (Agilent, USA), and Kapa HyperPrep Kits (Kapa Biosystems, USA) were used for constructing four DNA libraries. The prepared libraries with average fragment size from 451 to 461 bp were sequenced using an Illumina HiSeq 2500 sequencer with a HiSeq TruSeq Rapid Duo cBot sample loading kit and HiSeq Rapid PE cluster kit v2 (paired-end), yielding 108,689,404 reads for SDN1_day16; 96,381,152 reads for SDN2_day16; 92,719,398 for SPN1_day7; and 94,391,656 for SPN2_day7.

For metatranscriptomic sequencing, the quality of RNA (RNA integrity number from 7.5 to 9.4) was examined using the BioAnalyzer 2100 system with an RNA 6000 Nano Kit (Agilent, USA) before using the Ribo-Zero rRNA removal kit (Illumina) and TruSeq Stranded LT mRNA Library Prep kit v2 as described previously [23]. The four prepared libraries with average fragment size from 328 to 351 bp were sequenced using an Illumina HiSeq 2500 sequencer, yielding 97,486,966 reads for SDN1_day14; 92,150,650 for SDN2_day14; 94,830,584 for SPN1_day7; and 93,982,586 for SPN2_day7.

### Quality trimming, assembly, and gene prediction and binning of the metagenome

Raw reads with low quality (quality score <30) and short length (length< 36 bp) from four metagenomes—SDN1_day14, SDN2_day14, SPN1_day7, and SPN2_day7—were trimmed using Trimmomatic v0.39 [57]. The trimmed reads from all metagenomes were assembled into a single assembly by using Megahit v1.1.4 with the default setting [58]. Protein-coding genes were predicted using Prodigal 2.6.3 in the metagenome mode [59]. To recover genomes from the metagenome assembly, a binning algorithm—MaxBin 2.2.7—was applied [60]. The taxonomy affiliation of binned genomes was determined using the following steps. Firstly, the amino acid sequences of predicted protein-coding genes were mapped against the NCBI non-redundant protein database by using DIAMOND v0.9.26 [61] with a cutoff E value of 1 × e^−5^. Second, the closest taxonomic hit along with AAI of each predicted gene were extracted. Finally, the closest species were selected on the basis of the majority of hits, and AAIs were averaged to be the AAI between the binned genome and its closest species. CheckM v1.07 [62] was used to examine the completeness and contamination of binned genomes. Other information regarding binned genomes was assessed using QUAST 5.0.2 [63].

### Gene quantification and differential gene expression (DGE) analysis

After performing quality trimming by using Trimmomatic v0.39 [57], we mapped metatranscriptomic reads to the metagenome assembly by using Bowtie2 2.3.5.1 [64] and quantified them using featureCounts (under Subreads release1.6.4.). The read count table from featureCounts was applied to edgeR 3.26.7 [65] to analyze DGE between SDN and SPN communities with two biological replicates. Genes with log2-fold-change (log2 FC) value ≥ 2, a false discovery rate (FDR) ≤ 0.05, and an adjusted P < 0.05 were designated differentially expressed genes. The quantity of differentially expressed genes is presented as the count per million (CPM).

### Search for genes encoding MEHP transporter, DEHP/MEHP hydrolase, *o*-phthalic acid transporter, and phthaloyl-CoA decarboxylase

The hidden Markov model (HMM) was employed to identify the DEHP transporter and DEHP hydrolase involved in alkyl side-chains degradation of DEHP. The protein sequences of the DEHP hydrolase (NCU65476) of *Acidovorax* sp. strain 210-6 and MEHP transporter (WP_007297306.1) identified in *Rhodococcus jostii* RHA1 [66] were used for generating two HMM profiles. The TRAP transporter and UbiD-like phthaloyl-CoA decarboxylase involved in *o*-phthalic acid uptake and degradation were also searched using the HMM (http://hmmer.org v3.2.1) based on the six amino acid sequences of phthaloyl-CoA decarboxylase determined from nitrate-reducing and sulfate-reducing *o*-phthalic acid degraders [19, 35]. These HMMs were then used to search against the amino acid sequences of differentially expressed genes from SDN communities. InterPro [67] and SignalP 5.0 [68] webserver were applied to predict protein family and signal peptide of selective hydrolases, respectively.

### General gene annotation

The amino acid sequences of differentially expressed genes from SDN communities were annotated against the RefSeq non-redundant protein database (release 94) [69] using DIAMOND v0.9.26 with a cutoff E value of 1 × e ^−5^ [61] and the ortholog-based eggNOG mapper 5.0 [70]. The HMMER-based KofamKOALA [71] was employed to annotate binned genomes containing genes encoding DEHP hydrolase and phthaloyl-CoA decarboxylase that were identified in this study. β-oxidation genes–namely CoA transferase, acyl-CoA dehydrogenase, enoyl-CoA hydratase, and 3-hydroxyacyl-CoA dehydrogenase and thiolase–were selected if these genes were not an adjunct to any genes involved in aromatics degradation or amino acid metabolism. In addition, we used blastp in the NCBI or Uniprot database to annotate gene function manually.

### Isolation of DEHP-degrading Acidovorax sp. strain 210-6

On day 16, the DEHP-treated estuarine meocosm (∼20 mL) was transferred into a 250-mL serum bottle containing defined mineral minimal medium (200 mL) with DEHP (1 mM) as the sole carbon source and electron donor as well as sodium nitrate (10 mM) as the electron acceptor. The defined medium was prepared following an established protocol described previously [24]. After the degradation of almost all DEHP added to the medium, the culture was serially diluted (10^-1^ to 10^-7^), and transferred to other serum bottles with the same defined medium for further cultivation. The fourth subculture containing highly enriched DEHP-degrading denitrifier was spread on tryptone soy agar containing DEHP (1 mM) to obtain single colony. Most colonies were identified as *Acidovorax* spp. in the DEHP-containing plates through PCR by using the universal primers of the bacterial 16S rRNA gene (27F and 1492R) [48]. *Acidovorax* colonies were cultivated in the aforementioned defined medium to confirm their function. The isolate cultures were used to test their utilization of other DEHP-related substrates including MEHP, 2-ethyl-1-hexanol, *o*-phthalic acid, and benzoic acid.

### Genome sequencing, assembly, and annotation for *Acidovorax* sp. strain 210-6

The genomic DNA of *Acidovorax* sp. 210-6 was extracted using Prest Mini gDNA Bacteria Kit (Geneaid, Taiwan). The integrity of DNA was examined using Fragment Analyzer (Agilent, USA) showing the major peak size at 35kb. The PacBio shotgun library was constructed following the manufacturer’s multiplexed bacterial protocol for the SMRTbell Express TPK 2.0 kit (PacBio). Briefly, gDNA was sheared using Megaruptor 2 (Diagenode) to an average size of 8.9kb, purified, and condensed using Ampure. Single-stranded DNA was removed using nucleases provided in the kit. Fragments were then subjected to end-repair, A-tailing, and ligation to the adaptor of Barcoded Overhang Adapter kit 8A. DNA size of 6–20 kb was selected using the BluePippin gel cassette (Sage). The final library showed an average size of 8.5kb.

PacBio sequencing was carried out using the Sequel Sequencing Kit 3.0 with SMRT Cell 1M v3 LR and run on a Sequel sequencer, and the read file was generated from SMRTlink ICS v6.0. The *de novo* assembly was conducted using HGAP4.0 on SMRTlink v8.0 with a coverage depth of 258×, resulting in three polished contigs. The NCBI Prokaryotic Genome Annotation Pipeline was applied for genome annotation. The gene clusters of NCU65476 were visualized using Gene Graphics.

### Purification of extracellular DEHP/MEHP hydrolase from strain 210-6

The *Acidovorax* sp. 210-6 culture (2 L) grown on DEHP was centrifuged at 10,000 × g for 15 min to pellet down the bacterial cells, cell debris, and residual DEHP. Extracellular proteins (10 μg/mL; totally 700 mL) were filtered through a 0.22-μm nitrocellulose membrane (47-mm diameter; Millipore) and concentrated using the Amicon Ultra Centrifugal Filters to 7 mL (cut-off of 30 kDa). The resulting proteins were purified using the AKTA start purification system through DEAE Sepharose Fast Flow packed in the XK16 column (GE Healthcare, USA), followed by another purification using Phenyl Sepharose (GE Healthcare, USA).

### Proteomic analysis

For protein identification, active protein fractions were further separated through sodium dodecyl sulfate-polyacrylamide gel electrophoresis (SDS-PAGE; 4%–20% Bis-Tris Gel; GenScript, USA). Proteins in the gel slices were eluted in HEPES-K^+^ buffer (50 mM; pH 8.0) and trypsin-digested. The proteomic analysis was performed using an LC-nESI-Q Exactive MS model (Thermo Fisher Scientific, USA) coupled with an on-line nanoUHPLC (Dionex UltiMate 3000 Binary RSLCnano). Protein identification was performed using the Proteome Discoverer software (v1.4, Thermo Fisher Scientific) with SEQUEST search engine against all the protein sequences of the genome and the plasmids of *Acidovorax* sp. strain 210-6. All peptides were filtered with a q-value threshold of 0.01 (false discovery rate of 1%), and proteins were filter with a minimum of two peptides per protein, wherein only rank-1 peptides and the peptides in top-scored proteins were counted.

### DEHP and MEHP hydrolase activity assay

The hydrolase activities of protein fractions were determined in 50 mM HEPES-K^+^ buffer (pH 8.0) containing 0.2 mM of either DEHP or MEHP. The reaction was carried out at 30°C and sampled at 0 and 12 h. The remaining DEHP, MEHP, or *o*-phthalic acid in each reaction was extracted by ethyl acetate and detected using thin-layer chromatography and UPLC–APCI–HRMS.

### Phylogenetic analyses for phthaloyl-CoA decarboxylase and DEHP/MEHP hydrolase

Maximum likelihood trees were constructed in MEGAX [72] to elucidate the phylogeny of UbiD family decarboxylase and alpha/beta hydrolase. All amino acid sequences for this analysis were aligned without truncation by using MUSCLE [55] in MEGAX. The best amino acid substitution model for each tree was determined using the Model Test in MEGAX. The branch support was determined by bootstrapping 1,000 times

For the UbiD tree, apart from the amino acid sequences of *ubiD* in the *Acidovorax* sp. strain 210-6 genome and DGE in SDN metagenome, we selected 100 amino acid sequences of 3-octaprenyl-4-hydroxybenzoate carboxylase and four phenolic acid decarboxylase sequences (all manually annotated and reviewed) from the Uniprot database as references. One sequence of phthaloyl-CoA decarboxylase from *Azoarcus* Bin394 and eight sequences of phthaloyl-CoA decarboxylase from anaerobic phthalate degraders [19, 35], one sequence of iso-phthaloyl-CoA decarboxylase and its closely related sequences (phenolic acid decarboxylase subunit C) [74], one sequences of 2,5-furandicarboxylate decarboxylase, phenolic acid decarboxylase subunit C, and phenylphosphate carboxylase subunit alpha and beta were also included for this phylogenetic analysis. The UbiD maximum likelihood tree was constructed using the LG substitution model plus Gamma distribution rate.

The phylogeny of hydrolases responsible for the alkyl-side chain degradation on PAEs was also constructed. The amino acid sequences of alpha/beta hydrolases in the genome of *Acidovorax* sp. 210-6 and hydrolases derived from several aerobic *o*-phthalic acid-degrading *Actinobacteria* [14–16, 67, 75–77], *Alphaproteobacteria* (78, 79), *Firmicutes* [80], *Gammaproteobacteria* [81], and uncultured bacterium [82] were used for inferring maximum likelihood tree. The bacterial source, substrate specificity, and accession number of these aerobic hydrolases are listed in **Table S1**. We included the amino acid sequences of the most similar proteins to NCU65476 (identity ≥ 40%) from the NCBI and Uniprot database. The maximum likelihood tree was constructed under the WAG+F substitution model plus Gamma distribution rate.

### Data availability

The raw reads of 16S amplicon sequencing from SN, SBN, SDN, and SPN communities were deposited in Sequence Read Archive (SRA) of NCBI under accession PRJNA604667. The raw reads of the metagenome and metatranscriptome of SDN_day14 and SPN_day7 is available under the SRA experiments of PRJNA602375 in the NCBI. The strain 210-6 genome was deposited in the NCBI under accession GCA_010020825.1.

## Acknowledgment

This study was supported by the Ministry of Science and Technology of Taiwan (MOST 109-2221-E-001-002 and 110-2222-E-008-002) and Academia Sinica Career Development Award (AS-CDA-110-L13). Yi-Lung Chen is supported by Research Grants for New Teachers of College of Science, Soochow University, Taiwan. Po-Hsiang Wang is supported by the Research and Development Office and Research Center for Sustainable Environmental Technology, National Central University, Taiwan. We especially thank the High Throughput Genomics Core Facility hosted by the Biodiversity Research Center at Academia Sinica for conducting the next generation sequencing experiments. This core facility is funded by the Academia Sinica Core Facility and Innovative Instrument Project (AS-CFII-108-114). We also appreciate Yu-Ching Wu for operating UPLC– HRMS at the Small Molecule Metabolomics Core Facility, Institute of Plant and Microbial Biology.

